# First evaluation of contrast-enhanced micro-XCT as a tool for organismal and reproductive trait observations in corals based on scans of *Thesea nivea* Deichmann, 1936

**DOI:** 10.1101/2025.07.21.665969

**Authors:** Erin E. Easton

## Abstract

Micro-X-ray computed tomography (micro-XCT) with contrast enhancement is considered a non-destructive tool that is increasingly being applied to explore the internal and external structures of vertebrates and invertebrates. Although micro-XCT has been applied to corals to study skeletal features, contrast enhancement has not been applied to evaluate the utility for visualization of soft tissue structures in corals. This study is the first to evaluate the utility of contrast enhancement to visualize internal and external features, including soft tissue features, of the octocorals. Contrast staining of a *Thesea nivea* specimen with 2.5% Lugol’s iodine permitted visualization of soft tissues with sufficient density differences among tissues to distinguish skeletal elements, dermal layers, polyp internal structures, and oocyte stages and features. Thus, contrast-enhanced micro-XCT is a tool that could advance octocoral systematics. Further optimization of this approach could further enhance its utility for systematics and increase its efficiency through development of semi-automated pipelines for morphometric and histological analyses.

**Highlights:**

- Contrast-enhanced micro-XCT proves useful for octocoral systematics.
- Organismal and histological information for *T. nivea* was visualized with contrast.
- Sufficient density differences were observed to differentiate among soft tissues.

## 1. Introduction

X-ray computed tomography (XCT) has applications in numerous fields (reviewed in Withers et al., 2021), including metrology, manufacturing, engineering, food science, materials science, biomedical and life sciences, paleontology, and Earth sciences. It is considered a non-destructive tool (but see Faulwetter et al., 2013) to obtain information on internal and external structures at scales of meters to nanometers (Withers et al., 2021), depending on the resolution of the scanner. In the life sciences, micron-scale XCT (hereafter micro-XCT) has been used to study non-invasively the external and internal morphological characters of a polychaetes (Faulwetter et al., 2013), earthworms (Fernández et al., 2014), chick embryos (Metscher, 2009), frogs (Porro and Richards, 2017), and fish (Weinhardt et al., 2018). Compared to dissection-based analyses, application of micro-XCT for morphology-based systematics has advantages in not only being non-destructive but also in its utility for the rapid creation of high-resolution, three-dimensional, morphological data (Faulwetter et al., 2013) and histological information (Gutiérrez et al., 2018) that can be digitalized as virtual type material (i.e., cybertypes; see Buser et al., 2020; Faulwetter et al., 2013; Porro and Richards, 2017) that could serve for virtual dissections, biochemical modeling (Buser et al., 2020; Porro and Richards, 2017), and education (Buser et al., 2020; Porro and Richards, 2017).

These benefits are enhanced with the application of contrast stains to enhance visualization of soft tissues. With the application of chemical contrast stain, taxonomically informative morphological details of internal structures and histological information have been visualized in lumbricid earthworms (Fernández et al., 2014), crickets (Gutiérrez et al., 2018), frogs (Porro and Richards, 2017). Among the contrast stains used, Lugol’s iodine and other iodine contrast stains have shown promise for visualizing soft-tissue and skeletal structures, including delicate and small structures that are often difficult to visualize through traditional dissection methods, in a wide range of vertebrates and invertebrates (Fernández et al., 2014; Gutiérrez et al., 2018; Metscher, 2009; Porro and Richards, 2017). Despite the promise of contrast-enhanced micro-XCT for systematics, its application for systematics remains limited even for taxa, such as corals, that have been studied with XCT and micro-XCT for decades.

In corals, micro-XCT has been applied to study skeleton density (Duprey et al., 2012; Roche et al., 2010) and structural/morphological characteristics (Fabri et al., 2022; Kaandorp et al., 2005; Knackstedt et al., 2006; Kramer et al., 2022; Kruszyński et al., 2007), effects of ocean acidification on coral skeletons (Enochs et al., 2015; Hennige et al., 2020; Scucchia et al., 2021; Tambutté et al., 2015; Wolfram et al., 2022), coral surface area and volume relationships (Gutiérrez-Heredia et al., 2015; House et al., 2018; Laforsch et al., 2008), effects of turbulence and flow on coral skeletons (Iwasaki et al., 2018), growth patterns (Li et al., 2021; Li et al., 2020; Medellín-Maldonado et al., 2022; Yudelman and Slowey, 2022), effects of coral-symbiont relationships (Beuck et al., 2007), and model the effects of environmental or biological changes on coral skeletons (Wolfram et al., 2022). Most of these studies have been conducted on scleractinians corals, which have substantial calcium carbonate skeletons. In contrast, few studies have been conducted on octocorals. The first application of micro-XCT to octocorals was mapping of the stem canals of octocorals to understand their internal organization and their physiological and functional role in octocoral colony growth (Morales Pinzón et al., 2014).

Micro-XCT was also used to explore the effects of ocean acidification on the octocoral *Eunicea flexuosa* by Enochs et al. (2015). Urushihara et al. (2016) were the first to use micro-XCT to obtain biological and structural information for an octocoral, the white coral, *Corallium konojoi*. Early applications of micro-XCT on corals often report a limited or lack of detection of soft tissue as a methodological limitation using standard scanning or modifications such as phase-contrast and synchrotron micron-XCT (Morales Pinzón et al., 2014; Naumann et al., 2009; Urushihara et al., 2016). Despite not using contrast stain, Urushihara et al. (2016) were able to distinguish some information on the structures formed by living tissues, such as gastrovascular canals and polyp cavities, through application of edge-enhancement effect of synchrotron X-ray imaging.

To the best of my knowledge, no studies have used chemical contrast stains to visualize soft tissues in corals, so the utility of micro-XCT to explore internal soft tissues of corals has yet to be evaluated. Here, I conduct the first evaluation of the utility of contrast enhancement for corals using Lugol’s iodine and a specimen of the octocoral *Thesea nivea* Deichmann, 1936 to determine whether additional morphological characters are distinguishable compared to micro-XCT without contrast staining and thus whether this tool could be useful for visualization of organismal and reproductive traits, advancing octocoral systematics.

## 2. Materials and methods

### 2.1. Sample collection and preparation

A specimen of *Thesea nivea* (DFH33-536A) was collected by the remotely operated vehicle SubAtlantic Mohawk 18, operated by the UNCW Undersea Vehicles Program, on 29 September 2017 at 86 m depth at MacNeil Bank (28.0410 °N, 93.4933 °W). The specimen was preserved in 95% ethanol at -20°C. An ∼5 cm long piece of a terminal branch was clipped to include a terminal branching point and sent to the University of Texas High-Resolution X-ray Computed Tomography Facility (hereafter, UTCT). The raw sequencing data from Quattrini et al. (2024) is available from GenBank (SAMN38145133).

### 2.2. Micro-XCT imaging and processing

UTCT staff conducted a total of three scans on the specimen. The first scan was performed 16 December 2022 on the ∼5 cm (length) fragment (Fig. 1), unstained, with the NSI Helical scanner with the following parameters: Fein Focus High Power source, 90 kV, 0.15 mA, aluminum foil filter, Perkin Elmer detector, 0.25 pF gain, 2 fps, 1×1 binning, source to object 70.84 mm, source to detector 1053.225 mm, helical continuous CT scan, vertical extent 64 mm, pitch 6.39698 mm, 10 revolutions, 2.5 sets, no frames averaged, 0 skip frames, 15000 projections, 5 gain calibrations, 5 mm calibration phantom, data range [-5, 100] adjusted grayscale values, beam-hardening correction = 0.1. A total of 3697 slices of voxel size 13.5 µm were obtained. To increase resolution, the fragment was cut down to an ∼ 0.7 × 0.5 cm fragment at the branch junction (Fig. 1) and rescanned, unstained, on 28 February 2023 with the Zeiss Xradia 620 Versa with the following parameters: flat panel, 70kV, 8.5W, 0.1s acquisition time, 5 samples per view, detector 252.336 mm, source -14.249 mm, XYZ [-117, -1533, -203], camera bin 1, angles ±180, 3201 views, no filter, dithering, multi reference. The 16bit TIFF images were reconstructed by Xradia Reconstructor with center shift -0.735, beam hardening 0, theta 0, byte scaling [-0.002, 0.035], binning 1, and recon filter smooth (kernel size = 0.7). To explore the use of contrast enhancement to observe the soft tissues, the fragment used for the second scan was then stained in 2.5% Lugol’s for ∼27 h and was subsequently scanned on 22 August 2023 Zeiss Xradia 620 Versa with the following parameters: flat panel, 70kV, 8.5W, 0.09s acquisition time, 5 samples per view, detector 252.5 mm, source -14.3 mm, XYZ [468, -21729, 567], camera bin 1, angles ±180, 4501 views, no filter, dithering, multi reference. The 16bit TIFF images were reconstructed with center shift -0.839, beam hardening 0.2, theta 0, byte scaling [-0.02, 0.65], binning 1, and recon filter smooth (kernel size = 0.7). A total of 3697, 1470, and 1470 tomographic slices (archived at UTCT) were obtained respectively for scan 1, 2, and 3 with a voxel size 13.5 µm (scan 1) and 4.0 µm (scans 2 and 3).

**Fig. 1.**
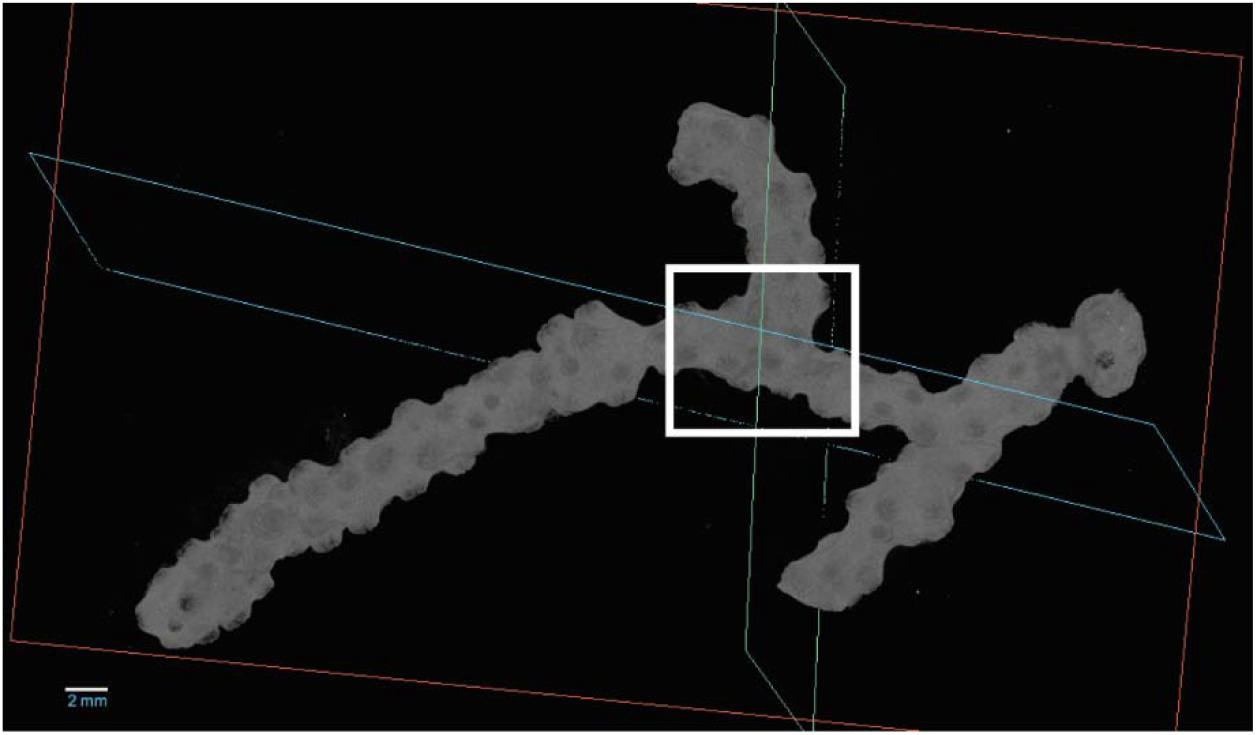
3D rendering of unstained micro-XCT scan of a ∼5 cm long fragment of *Thesea nivea* specimen DFH33-536A with a white box indicating the approximate boundaries of the ∼0.7 × 0.5 cm fragment subsequently scanned with and without contrast (Fig. 2). Scanned at The University of Texas High-Resolution X-ray Computed Tomography Facility (UTCT) with the NSI Helical scanner. Voxel size = 13.5 µm. Rendered in Dragonfly 2022.2 Build 1409.

Dragonfly 2022.2 Build 1409 (Comet Technologies Canada, Inc., Montreal, Canada; software available at https://www.theobjects.com/dragonfly) was used for three-dimensional reconstructions and analyses, including measurements of characters, of the tomographic images. In DragonFly, a threshold of 10500 (first scan; Fig. 1), 7500 (second scan, Fig. 2A-D) and 6000/5000 (3D/2D, third scan, Fig. 2E-H) were used to segment and remove the bulk of the background noise prior to viewing 3D reconstructions and exporting images; additional background noise pixels were removed with Dragonfly ROI define range and ROI painter 3D freehand tool and overwritten to a value of 0. Octocoral morphological terminology follows Bayer et al. (1983) and (Fabricius and Alderslade, 2001).

**Fig. 2.**
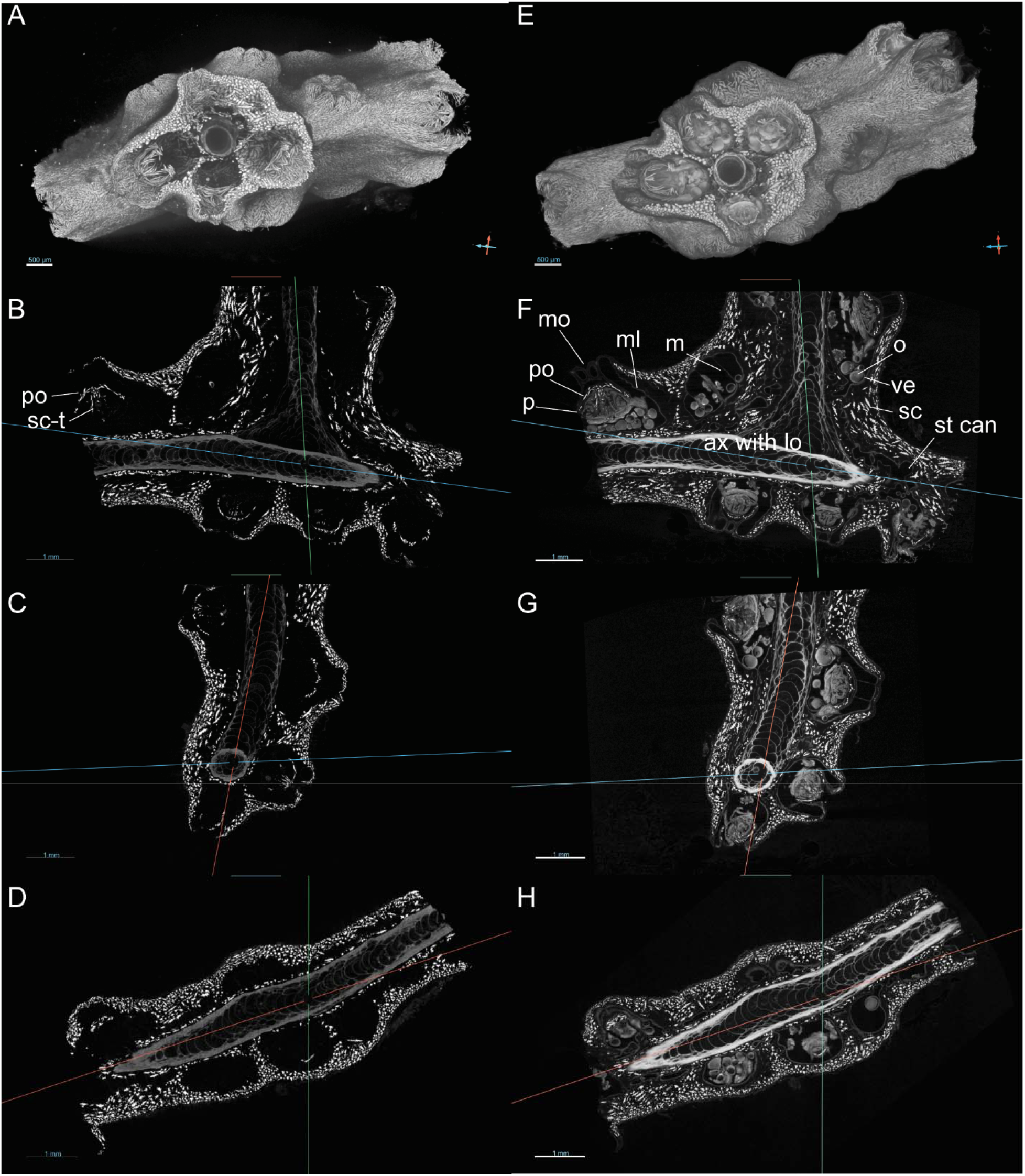
3D renderings and approximately equal views on three planes of an ∼0.7 × 0.5 cm fragment of *T. nivea*, showing the skeletal elements visualized by scanning without contrast (A-D) and the additional morphological features visible with contrast enhancement with ∼27 h of staining with 2.5% Lugol’s iodine (E-H). ax = axis, lo = loculus, m = mesenteries, ml = mesenterial lining, mo = mouth opening, o = oocyte, p = polyp, po = point, sc = sclerite, sc-t = sclerites in tentacles, st can = stem canals, and ve = vitelline envelope. Morphological terminology follows Bayer et al. (1983) and (Fabricius and Alderslade, 2001). Scanned with a Zeiss Xradia 620 Versa. Voxel size = 4.0 µm. Rendered in Dragonfly 2022.2 Build 1409.

## 3. Results

Thresholds used to isolate the specimen from air differed between scans with the lowest thresholds (5000/6000) used for the contrast-enhanced scan. Without contrast, the axis and sclerites were visualized but visualization of soft tissues was not achieved. Contrast enhancement allowed visualization of soft tissues and sufficient differentiation of soft tissues to identify internal and external morphological features and histological information; however, the threshold between air and stained coral tissues was not sufficiently distinct to completely distinguish between air and some soft tissues. Skeletal axis with loculi and a central core (Fig. 2–3), polyp (autozooid) cavities, and sclerites were visible with and without staining (Fig. 2). The arrangement of the coenenchyme and polyp sclerites are visible externally and internally.

Sclerites are observed along the central axis and throughout the coenenchyme with greater apparent densities near the surface than between polyps, and along the axis. Without contrast staining (Fig. 2 A-D), tissues were largely not distinguishable from air, so polyp cavities were visible but the boundaries with the surrounding tissues were not distinguishable. In contrast, Lugol’s iodine sufficiently stained the sample to allow for visualization of soft tissues (Fig. 2 E-H), including polyps, dermal layers, and gametes, as well as the skeletal elements. Autozooid mean width (1.55 ± 0.29 mm [sd]) was considerably larger (∼26%) than measured with contrast (1.23 ± 0.18 mm, n = 7) and polyp mean height (0.53 ± 0.04 mm) was considerably lower (25%) than measured with contrast (0.71 ± 0.08 mm, n = 3, respectively). Estimated autozooid mean heights and polyp mean widths were comparable between scans with contrast (1.23 ± 0.12 mm and 0.91 ± 0.19 mm, respectively; n = 7) and without contrast (1.22 ± 0.11 mm and 0.86 ± 0.13 mm, respectively; n = 3). Sufficient detail was observed to see the mesenterial lining of the polyp cavity, oocytes separated by mesenteries, the arrangement of point and collar sclerites relative to soft tissues, the mouth opening, stem canals and their linings, and oocyte characteristics with notable density differences. At least two cohorts of oocytes are visible with early stage, previtellogenic oocytes being adjacent to late-stage, vitellogenic oocytes that are ready to spawn. The nucleus, nucleolus, yolk, and vitelline envelope are visible (Fig. 3).

**Fig. 3.**
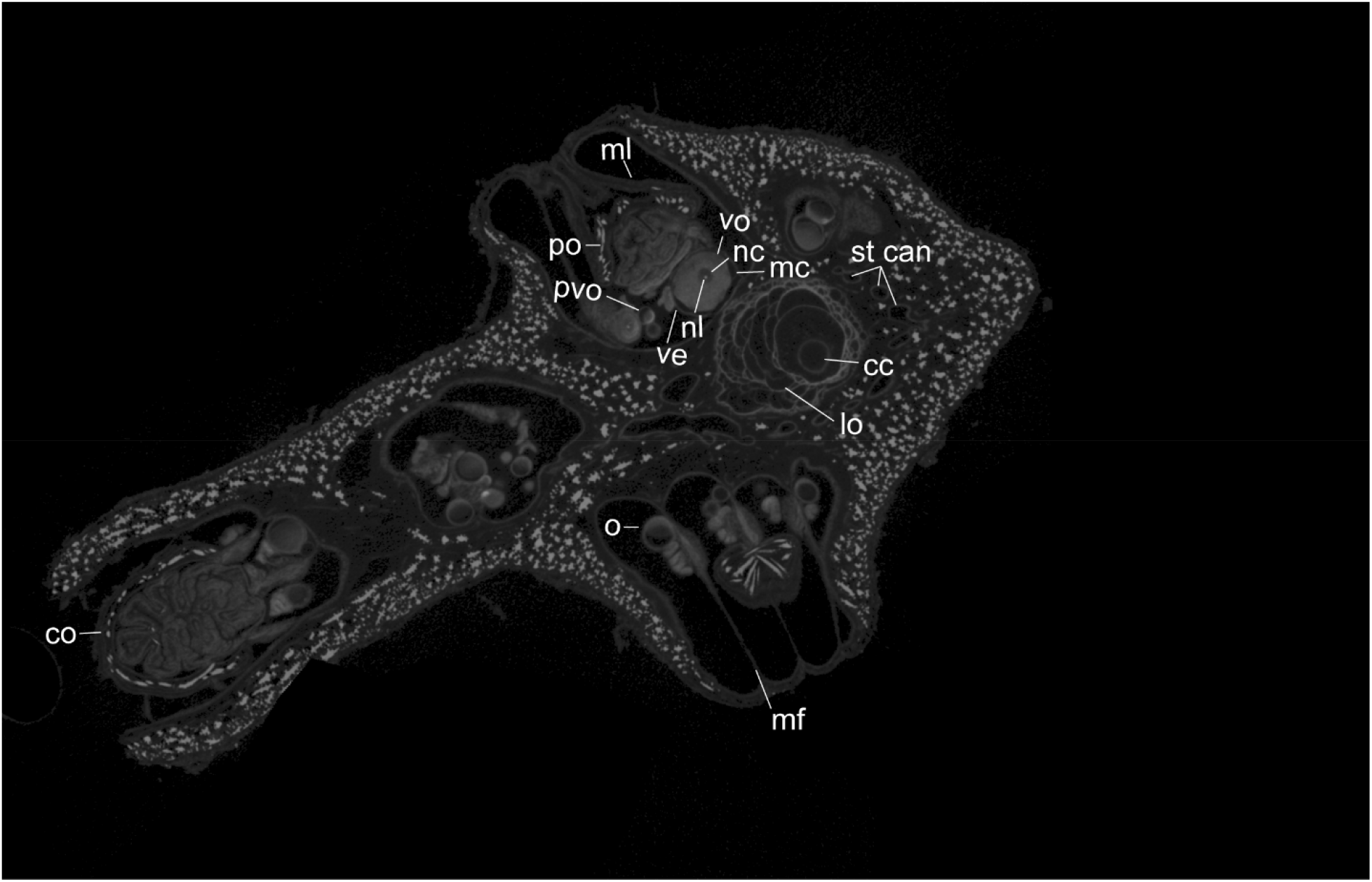
Micro-XCT slice of *T. nivea*, showing some of the additional histological features visible with contrast enhancement with ∼27 h of staining with 2.5% Lugol’s iodine. cc = central canal, co = collar, lo = loculus, mc = mesoglea capsule, mf = mesentery filament, ml = mesenterial lining, nc = nucleus, nl = nucleolus, o = oocyte, po = point, pvo = previtellogenic oocyte, st can = stem canals, ve = vitelline envelope, and vo = vitellogenic oocyte. Morphological terminology follows Bayer et al. (1983) and (Fabricius and Alderslade, 2001). Scanned with a Zeiss Xradia 620 Versa. Voxel size = 4.0 µm. Rendered in Dragonfly 2022.2 Build 1409.

## 4. Discussion

Micro-XCT without contrast revealed skeletal features of the octocoral *T. nivea* but soft tissues were largely indistinguishable as noted by other studies (Morales Pinzón et al., 2014; Naumann et al., 2009; Urushihara et al., 2016). In contrast, micro-XCT with contrast enhancement permitted visualization of soft tissues with sufficient density differences and detail to differentiate among oocyte stages and characteristics (i.e., nucleus, nucleolus, yolk, and vitelline envelope), among soft tissue layers (i.e., mesoderm and gastrodermis) to better visualize the internal structures, and the arrangement of skeletal elements internally and externally in *T. nivea*. Although sclerites in the coenenchyme and polyp serve as good markers for polyp width and autozooid cavity height because of their location near the mesenterial linings, contrast-staining allowed for more accurate measurements of polyp and autozooid cavity dimensions, in particular for the autozooid cavity width and polyp height measurements. As expected, higher resolution was achieved with a smaller region of interest as maximum resolution is related to the object’s dimensions (Withers et al., 2021). The goal of this study was not to optimize techniques but to evaluate whether contrast-enhanced micro-XCT could be used as a tool for visualization of organismal and reproductive traits in octocorals, thus enhancing octocoral systematics by obtaining additional morphometric and histological information for octocorals compared to micro-XCT without contrast staining.

Here, I demonstrated that 2.5% Lugol’s iodine was an effective medium for visualizing not only skeletal features visible without contrast enhancement but also internal soft tissues essential for describing organismal and reproductive traits in *T. nivea*. I acknowledge that this evaluation was applied to a single species and the utility to visualize the reproductive and organismal traits comprised of soft tissues may differ by taxon, but this application and methods presented herein provide a baseline for its application in octocoral systematics, ecology, and functional studies. Such future applications may require optimizations to the approach, so I will shift discussion to the potential next steps to optimize this approach for octocoral systematics.

Although the methods presented here are a first application of contrast-enhanced micro-XCT approaches in octocorals, detail and distinction among soft tissue features may be possible with modifications to the scanning protocol, including using a nano-XCT, using a scanner with helical scanning modes, optimizing contrast enhancement protocols, using density calibration techniques and applying image improvement protocols to improve ability to autosegment tissues (Chalker et al., 1985; Duprey et al., 2012; Faulwetter et al., 2013; Laforsch et al., 2008; Roche et al., 2010), and testing application for image analysis for thresholding, segmentation, and quantification of morphometrics.

Nano-scale XCTs and helical scanning features are worth exploring to achieve greater resolution and visualization of smaller characteristics; however, there are tradeoffs between sample size and resolution as well as scan time and thus cost to achieve higher resolution (see Withers et al., 2021). Further, higher resolution scans and higher-energy scans would increase the X-ray dose, in turn, increasing the potential destructive effects of XCT (Withers et al., 2021). For example, higher dosages could result in damage to DNA and compromise the integrity of the sample for future molecular analyses, which could be compounded by effects of the contrast stain used and the condition of the sample used. Although micro-XCT is largely considered non-destructive and non-invasive, the application of stain inherently makes the process invasive, even if minimally so, because stains are often not fully washed out (Fernández et al., 2014).

Nevertheless, contrast-enhanced micro-XCT was successfully applied to decades old museum specimens of earthworms (Fernández et al., 2014) demonstrates that this method could likewise be applied to museum collections of octocorals. Because the effects of micro-XCT and contrast stains on molecular material has revealed mixed results (see Faulwetter et al., 2013) and their effect on molecular material of preserved octocorals have not been explored, this method would need to be tested on museum collections to determine whether there are negative effects on specimens and if there is an optimal methodology or processing order to minimize effects and maximize recoverable information from type specimens. XCT has also been conducted on living scleractinians (Laforsch et al., 2008), so application of this method, with or without, contrast stains could also be explored to evaluate its utility and the potential destructive effects for living octocorals in aquaculture or after live collections prior to more destructive studies.

Further testing may also reveal optimal stains and staining protocols for histological information and autosegmentation. Lugol’s iodine was selected for this test because mixtures of iodine and potassium iodine have been identified as optimal stains for differentiation of tissues, are largely considered non-destructive, and will leach out of samples (see, e.g., Fernández et al., 2014; Gutiérrez et al., 2018; Metscher, 2009; Weinhardt et al., 2018). In addition, phosphotungstic acid (PTA) staining may be a viable option because, like Lugol’s iodine, it provided a strong contrast for differentiating chick embryo (Metscher, 2009), earthworm (Fernández et al., 2014), and teleost fish tissues (Weinhardt et al., 2018). Lugol’s iodine precipitates can decrease its dynamic range for tissue analysis (Weinhardt et al., 2018) and PTA is acidic and can take longer to penetrate than iodine solutions (Metscher, 2009), so decalcification could be an issue in corals. Different stains, stain concentration, and medias for preservation (e.g., dry, ethanol, formalin), staining (ethanol, methanol, formalin, water), and scanning may have an effect on tissue shrinkage and stain absorption (see, e.g., Faulwetter et al., 2013; Gutiérrez et al., 2018; Metscher, 2009; Weinhardt et al., 2018). Dual or multiple stains may be appropriate for optimizing histological information obtained from contrast-enhanced micro-XCT. Stains may also prove useful for exploring effects on calcium carbonate skeletal elements if they differentially stain these elements. Pairing contrast-stain methods with scanning electron microscopy (SEM) and chemical and histological analyses could further contribute to our understanding of coral morphology and physiology. Pairing these methods could also further test the utility of micro-XCT to differentiate characteristics and to elucidate additional taxonomic and ecological information (e.g., Beuck et al., 2007), such as chitinous features if XCT methods could differentiate such chitinous features (Vandepas et al., 2023).

Automated pipelines for segmentation and morphometrics would improve the utility of contrast staining and micro-XCT. Such pipelines have been developed for other fauna, e.g., teleost fish (Weinhardt et al., 2018) and foraminiferal-algal nodule samples (Chandra et al., 2024). Some automated pipelines have been developed for specific tasks in corals, e.g., for analyzing indeterminate growth forms in the scleractinian *Madracis mirabilis* (Kruszyński et al., 2007) and for semi-automated stem canal tracking in the octocoral *Muricea muricata* (Morales Pinzón et al., 2014). These pipelines could be modified and integrated with pipelines for contrast-stained samples to optimize them for automated segmentation and morphometric analyses for integrative systematics and revisionary taxonomy, including for application to type specimens that can only be analyzed with nondestructive methodologies. For example, applications and pipelines could be developed to test whether volumes, areas, and quantities of tissues, sclerites, canal, axis, and other features could be calculated using automated, semi-automated, and artificial intelligence approaches. The raw data can be archived as done with this dataset at the UTCT and that data and the processed outputs can be made available in public repositories for subsequent analyses, virtual dissection, digital dissection, and educational purposed (see Buser et al., 2020 for Workflows and review of repositories and outputs).

## 5. Conclusions

Contrast-enhanced micro-XCT provided additional organismal and histological information for *T. nivea* compared to micro-XCT without contrast enhancement. This method permitted visualization of soft tissues with sufficient density differences among tissues to different mesoderm and gastrodermis layers, different stages and features of oocytes, as well as the arrangement of skeletal elements internally and externally. Thus, this tool shows promise for octocoral systematics, ecological, and functional studies. The utility of the tool could be enhanced by further testing to optimize protocols, e.g., to test different contrast staining protocols and image analysis protocols to facilitate a semi-automated approach, to collect morphometric and histological information. Its utility would be further enhanced with integration with traditional techniques for a more comprehensive systematics approach to aid in the description and revision of octocoral taxa. Such optimizations will result in enhanced biological and ecological information for octocorals and provide cybertypes for subsequent revisions, virtual dissection, and education.

## Acknowledgements

Research cruise and ROV operations funding provided by NOAA’s Deep-Sea Coral Research and Technology Program, Flower Garden Banks National Marine Sanctuary, and National Marine Sanctuary Foundation. Funding for micro-XCT imaging was supported by UTRGV CAREER development support to the author and NSF Division of Earth Science Instrumentation and Facilities Program (NSF EAR-2223808) and NASA (80NSSC23K0199) awards to UTCT. This publication was made possible by the National Oceanic and Atmospheric Administration, Office of Education Educational Partnership Program award NA16SEC4810009 and NA21SEC4810004. Its contents are solely the responsibility of the award recipient and do not necessarily represent the official views of the U.S. Department of Commerce, National Oceanic and Atmospheric Administration. I would like to thank J. Maisano for scanning support and A. Rossin for reviewing imagery and confirming reproductive stages. I thank members of my laboratory and the two anonymous reviewers whose comments helped improve this article.

